# Development of monoclonal antibody-based blocking ELISA for detecting SARS-CoV-2 exposure in animals

**DOI:** 10.1101/2023.03.11.532204

**Authors:** Fangfeng Yuan, Chi Chen, Lina M. Covaleda, Mathias Martins, Jennifer M. Reinhart, Drew R. Sullivan, Diego G. Diel, Ying Fang

**Author notes:** These authors contributed equally (Fangfeng Yuan, Chi Chen). To whom correspondence should be addressed: (Ying Fang).

## Abstract

The global pandemic of severe acute respiratory syndrome coronavirus 2 (SARS-CoV-2) poses a significant threat to public health. Besides humans, SARS-CoV-2 can infect several animal species. Highly sensitive and specific diagnostic reagents and assays are urgently needed for rapid detection and implementation of strategies for prevention and control of the infection in animals. In this study, we initially developed a panel of monoclonal antibodies (mAbs) against SARS-CoV-2 nucleocapsid (N) protein. To detect SARS-CoV-2 antibodies in a broad spectrum of animal species, a mAb-based bELISA was developed. Test validation using a set of animal serum samples with known infection status obtained an optimal percentage of inhibition (PI) cut-off value of 17.6% with diagnostic sensitivity of 97.8% and diagnostic specificity of 98.9%. The assay demonstrates high repeatability as determined by a low coefficient of variation (7.23%, 6.95%, and 5.15%) between-runs, within-run, and within-plate, respectively. Testing of samples collected over time from experimentally infected cats showed that the bELISA was able to detect seroconversion as early as 7 days post-infection. Subsequently, the bELISA was applied for testing pet animals with COVID-19-like symptoms and specific antibody responses were detected in two dogs. The panel of mAbs generated in this study provides a valuable tool for SARS-CoV-2 diagnostics and research. The mAb-based bELISA provides a serological test in aid of COVID-19 surveillance in animals.

**IMPORTANCE:** Antibody tests are commonly used as a diagnostic tool for detecting host immune response following infection. Serology (antibody) tests complement nucleic acid assays by providing a history of virus exposure, no matter symptoms developed from infection or the infection was asymptomatic. Serology tests for COVID-19 are in high demand, especially when the vaccines become available. They are important to determine the prevalence of the viral infection in a population and identify individuals who have been infected or vaccinated. ELISA is a simple and practically reliable serological test, which allows high-throughput implementation in surveillance studies. Several COVID-19 ELISA kits are available. However, they are mostly designed for human samples and species-specific secondary antibody is required for indirect ELISA format. This paper describes the development of an all species applicable monoclonal antibody (mAb)-based blocking ELISA to facilitate the detection and surveillance of COVID-19 in animals.

## INTRODUCTION

The causative agent of Coronavirus Disease 2019 (COVID-19), severe acute respiratory syndrome-related coronavirus 2 (SARS-CoV-2) is a new member of the family *coronaviridae* within the order *Nidovirales* (1). Nidoviruses are a group of positive-stranded RNA viruses, which replicate through a nested 3’-co-terminal set of subgenomic mRNAs, each possessing a common leader and a poly-A tail (2). The coronaviruses have an intriguing distant evolutionary relationship to other members of the order *Nidovirales,* but possess unique characteristics among currently known +RNA viruses. The coronavirus virion has a characteristic crown-like appearance with spike (S), membrane (M) and envelope (E) proteins inserted into the phospholipid-bilayered envelope. Inside the lipid bilayers, the RNA genome is packaged with a nucleocapsid (N) composed of N proteins. The replicase-associated genes, ORF1a and ORF1b, situated at the 5’-end of the viral genome. They encode two large polyproteins, pp1a and pp1ab, which are cleaved by viral encoded proteases to generate 16 known functional nonstructural proteins (nsp 1-16). The 3’-end of the viral genome encodes four major structural proteins: S, M, E and N proteins, and several other minor structural and accessory proteins (3). Host antibody responses induced by SARS-CoV-2 infection are mainly directed against S and N proteins (4).

SARS-CoV-2 has a broad host range (5). Besides humans, SARS-CoV-2 has been reported to infect multiple animal species, including cat (6), tiger (7), lion (7), snow leopard (8), deer (9), mink (10), dog (11, 12), etc. These findings cause great concerns on the potential for human to animal and animal to human transmission, along with the appearance of viral mutations as the virus spillover between species. Highly sensitive and specific diagnostic reagents and assays are urgently needed for rapid detection and implementation of control and prevention strategies.

Current diagnostic assays for SARS-CoV-2 detection mainly target viral nucleic acids or host antibodies against the viral infection. Nucleic acid tests detect active virus replication and shedding, while antibody tests reveal the previous exposure to the virus (13, 14). The fact that SARS-CoV-2 is capable of infecting a diverse range of animal species causes challenges for antibody test development, as certain reagents such as species-specific secondary antibodies are not commercially available for most animal species. Neutralization tests are an option to screen all animal species for SARS-CoV-2 neutralizing antibodies. However, it has limitations for large-scale field surveillance (15, 16). In contrast to the traditional indirect Enzyme-Linked Immunosorbent Assay (iELISA), monoclonal antibody (mAb)-based blocking ELISA (bELISA) is capable of detecting host antibodies independent of species-specific secondary antibody reagents (17). The bELISA was reported to be able to provide similar level of sensitivity as traditional indirect ELISAs, but with higher level of specificity (18). In this study, a panel of mAbs against SARS-CoV-2 N protein was generated, and a mAb #127-3-based bELISA was developed. Subsequently, the bELISA was applied to detect seroconversion in an experimental cat infection study (19) and diagnosis of SARS-CoV-2 specific antibody response in dogs from a pet animal clinic.

## RESULTS

### Generation and characterization of mAbs against SARS-CoV-2 N protein

To produce N antigen for mice immunization, synthetic gene of SARS-CoV-2 Wuhan-hu-1 strain was cloned and expressed as a His-tagged recombinant protein. On SDS-PAGE analysis, the purified N protein showed a single band with predicted molecular mass around 47.4 kDa (Figure 1A). The identity of the recombinant N protein was further confirmed on western blot using anti-His-tag antibody (Figure 1A).

**Figure 1.**
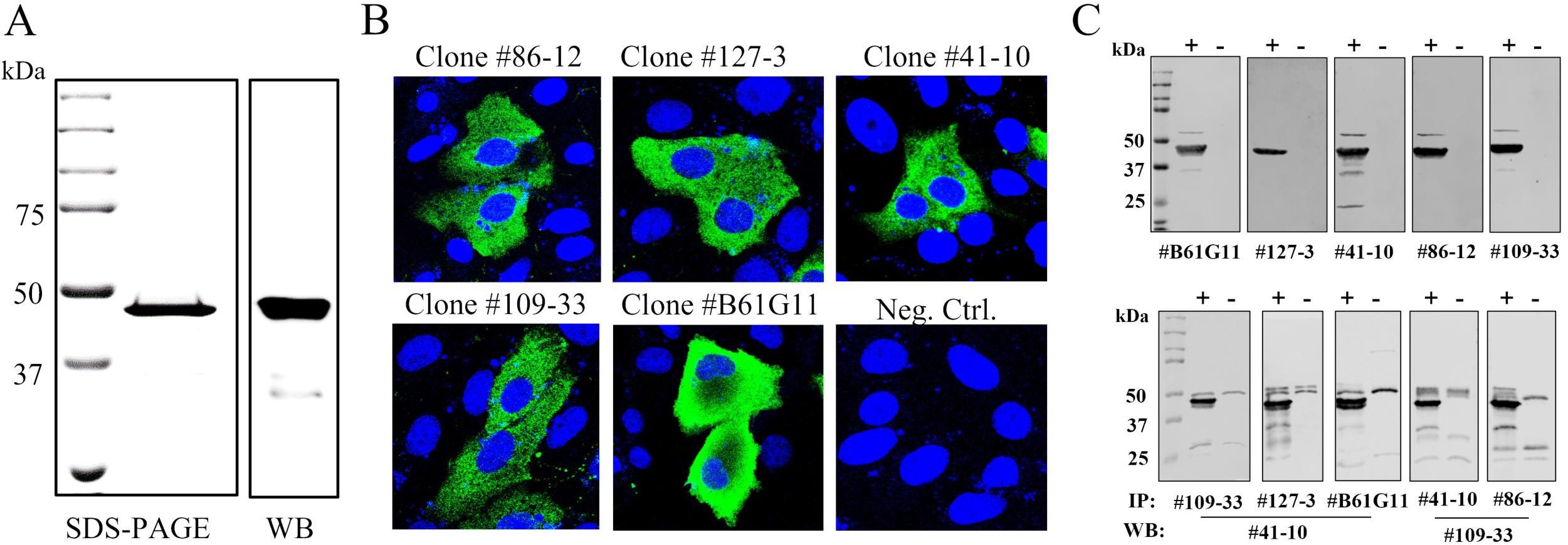
SARS-CoV-2 N antigen preparation and mAb characterization. **(A)** Recombinant N antigen expression and detection. Left panel, SDS-PAGE gel electrophoresis of recombinant N protein, followed by Coomassie blue staining; right panel, Western blot detection of His-tagged N protein. The membrane was stained with anti-His tag antibody. **(B)** IFA detection of the N protein expressed in transfected MARC-145 cells. Fixed cells were stained by the corresponding mAb and FITC-conjugated goat anti-mouse IgG was used as the secondary antibody. Nuclei were counterstained with DAPI (blue). **(C)** MAb reactivity tested on Western blot and immunoprecipitation (IP). Lysates from transfected 293T cells expressing the N protein were harvested and utilized for WB and IP analysis.

To generate the SARS-CoV-2 specific mAbs, mice were immunized with N antigen. After the fusion of mice splenocytes with myeloma cells, supernatants from the resulting hybridoma cells were screened by IFA using transfected MARC-145 cells expressing N protein. A total of 4 mAbs (clone #41-10, 86-12, 109-33, 127-3) were obtained. One additional mAb B6G11 previously developed in Diel’s lab (20) was included in the analysis. IFA result showed that all 5 mAbs recognized N proteins expressed in MARC-145 cells (Figure 1B). Using the cell lysate of transfected 293T cells that express N protein, this panel of mAbs was determined to be able to detect the N protein by western blot and immunoprecipitation (IP) (Figure 1C). To further determine if this panel of mAbs recognizes the N protein in virus-infected cells, Vero cells infected with SARS-CoV-2 variants, including B.1, WA1, P.1, B.1.1.7, and B.1.617.2, were subjected to IFA. The results showed that this panel of mAbs had different levels of reactivity with each of the variant, of which the mAb #127-3 and B61G11 had strong reactivity, #41-10 and #86-12 had moderate reactivity, while #109-33 had weak reactivity (Table 1).

**Table1.** Reactivity of mAbs with different SARS-CoV-2 variants.

| mAb clone# | B.1 | WA1 | P.1 | B.1.1.7 | B.1.617.2 |
| --- | --- | --- | --- | --- | --- |
| 41-10 | ++ | ++ | ++ | ++ | ++ |
| 86-12 | ++ | ++ | ++ | ++ | ++ |
| 109-33 | + | + | + | + | + |
| 127-3 | +++ | +++ | +++ | +++ | +++ |
| B6G11 | +++ | +++ | +++ | +++ | +++ |

“+” weak reactivity, “++” moderate reactivity, “+++” strong reactivity.

The mAb cross-reactivity with other common coronaviruses was further evaluated. We tested N proteins of common coronaviruses from SARS-CoV-2 susceptible host species, including the four human coronaviruses, two feline coronaviruses, two canine coronaviruses, mink and ferret coronaviruses (Table 2). Flag-tagged N proteins from each of these viruses were expressed in transfected cells. IFA results showed that mAb #86-12 can cross-react with the N protein of SARS-CoV, HCoV-OC43, and CCoV-Type 1, while mAb #B61G11 can cross-react with the N protein of SARS-CoV. In contrast, mAb #41-10, #109-33, and #127-3 did not cross-react with any of the N proteins from corresponding coronaviruses.

**Table 2.**
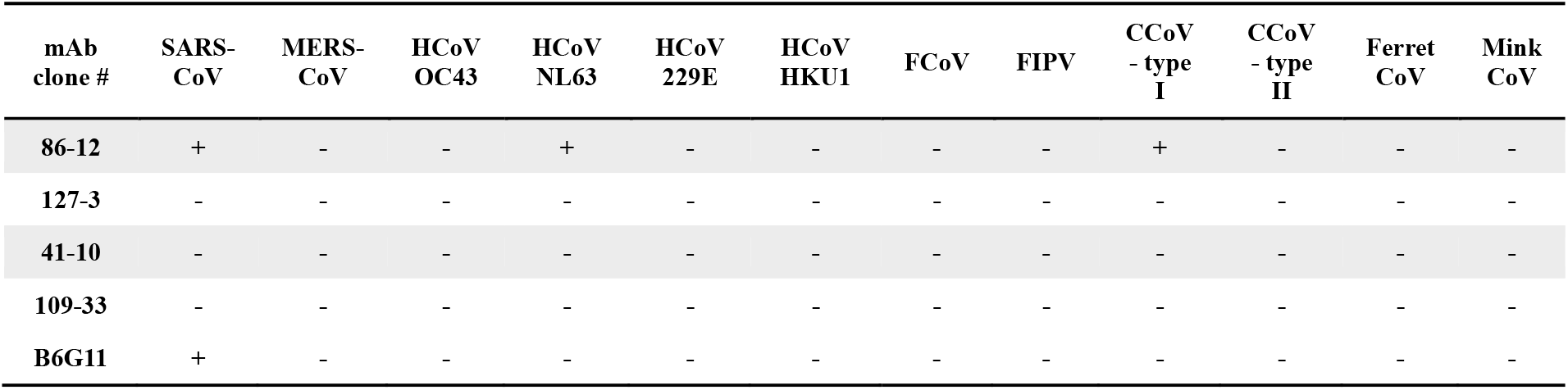
Cross-reactivity of mAbs with other coronaviruses.

| mAb clone # | SARS-CoV | MERS-CoV | HCoV OC43 | HCoV NL63 | HCoV 229E | HCoV HKU1 | FCoV | FIPV | CCoV - type I | CCoV - type II | Ferret CoV | Mink CoV |
| --- | --- | --- | --- | --- | --- | --- | --- | --- | --- | --- | --- | --- |
| 86-12 | + | - | - | + | - | - | - | - | + | - | - | - |
| 127-3 | - | - | - | - | - | - | - | - | - | - | - | - |
| 41-10 | - | - | - | - | - | - | - | - | - | - | - | - |
| 109-33 | - | - | - | - | - | - | - | - | - | - | - | - |
| B6G11 | + | - | - | - | - | - | - | - | - | - | - | - |

### Development and validation of #127-3 mAb-based bELISA

In order to detect anti-N antibody response in multiple animal species (independent of species-specific reagents), we further developed a mAb-based bELISA. Since mAb #127-3 had strong reactivity with different SARS-CoV-2 variants, and this mAb did not cross react with the other common coronaviruses and SARS-CoV-1, mAb #127-3 was selected for the assay development.

#### Establishment of serum standards

Initially, a set of internal control serum standards were established using cat sera collected from our previous study (19). A group of 24 cats were experimentally infected with SARS-CoV-2 virus (D614G, Delta, and Omicron). Serum samples collected from the cats at 14 days post infection were pooled into a single lot of positive control serum. Similarly, large quantities of the known negative cat sera was pooled into a single lot of negative control serum. The positive control standards were set as three levels in the indirect ELISA, including high-positive (OD of 1.5-2.0), medium-positive (OD of 1.0-1.5), and low-positive (OD of 0.8-1.0), while the negative control standard generated an OD of less than 0.3 in the indirect ELISA (Figure 2A). Using the positive and negative control standards, bELISA conditions were optimized by checkerboard titration of the antigen (N protein), biotinylated mAb #127-3, HRP-conjugated streptavidin, blocking and sample buffer component, incubation temperature and time, etc. With the optimized test conditions, the bELISA generated percentage of inhibition (PI) value 75-85% for high-positive standard, 55-65% for medium-positive standard, 35-45% for low-positive standard, and approximate 0% for negative control standard (Figure 2B).

**Figure 2.**
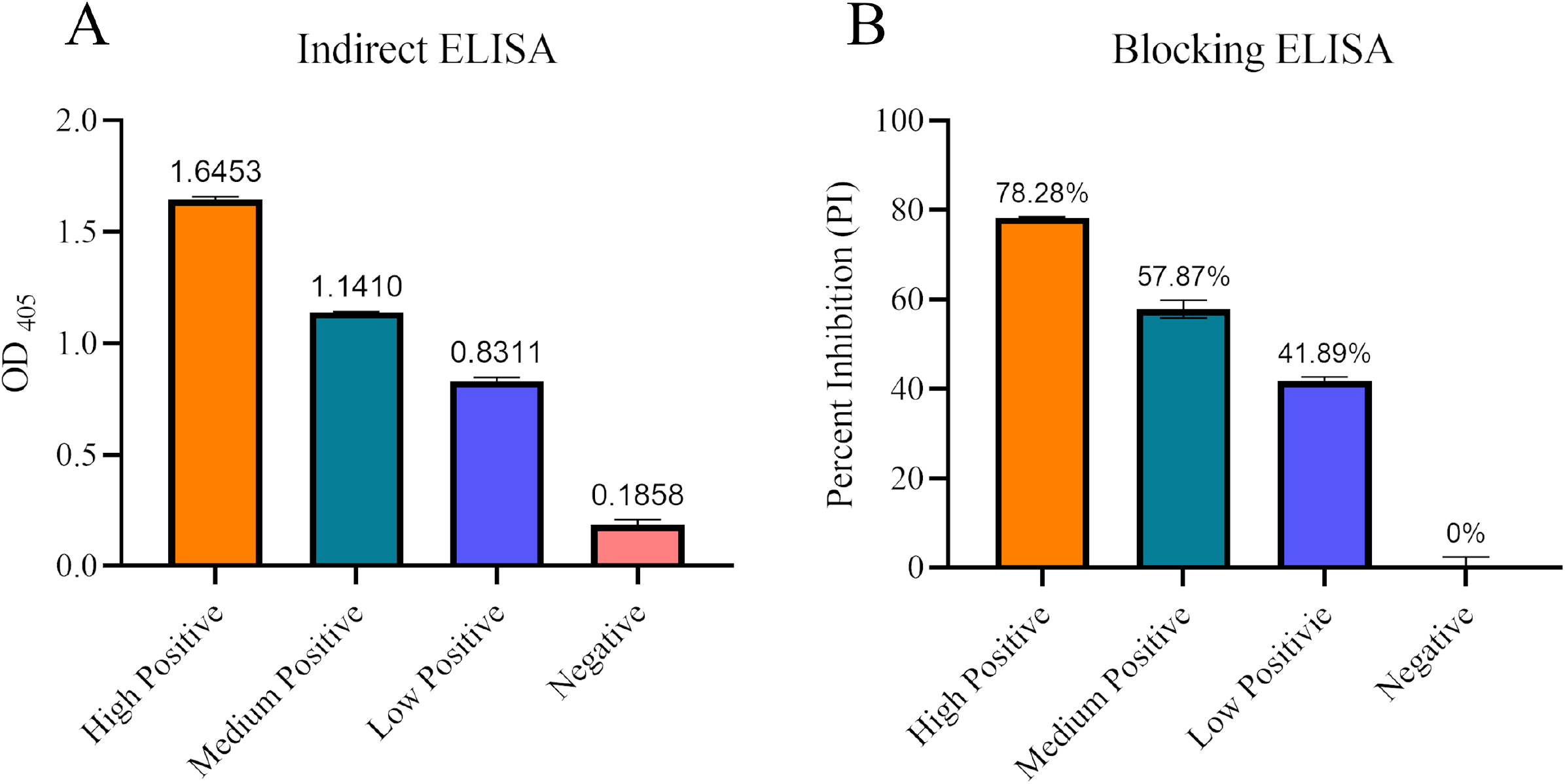
Establish positive and negative control standards. A set of internal control serum standards was prepared using experimental cat serum and assayed by indirect ELISA (**A**) and blocking ELISA (**B**). X-axis represents the positive and negative controls. Y-axis shows the OD_405_ for indirect ELISA and PI for bELISA. Each control standard was highlighted in different colors and mean value was displayed on top of each column.

#### Analytical sensitivity of bELISA

Analytical sensitivity of the bELISA was determined by using the high-positive and negative control standards. Standard sera were titrated with two-fold serial dilutions in triplicate. As shown in Figure 3, a dilution of 1:128 was the highest dilution that generates a statistical difference (p < 0.01) between the positive and negative control standards. A 1:4 dilution of the sample was selected for the bELISA, as it maximized the discrimination between positive and negative results and minimized background interference.

**Figure 3.**
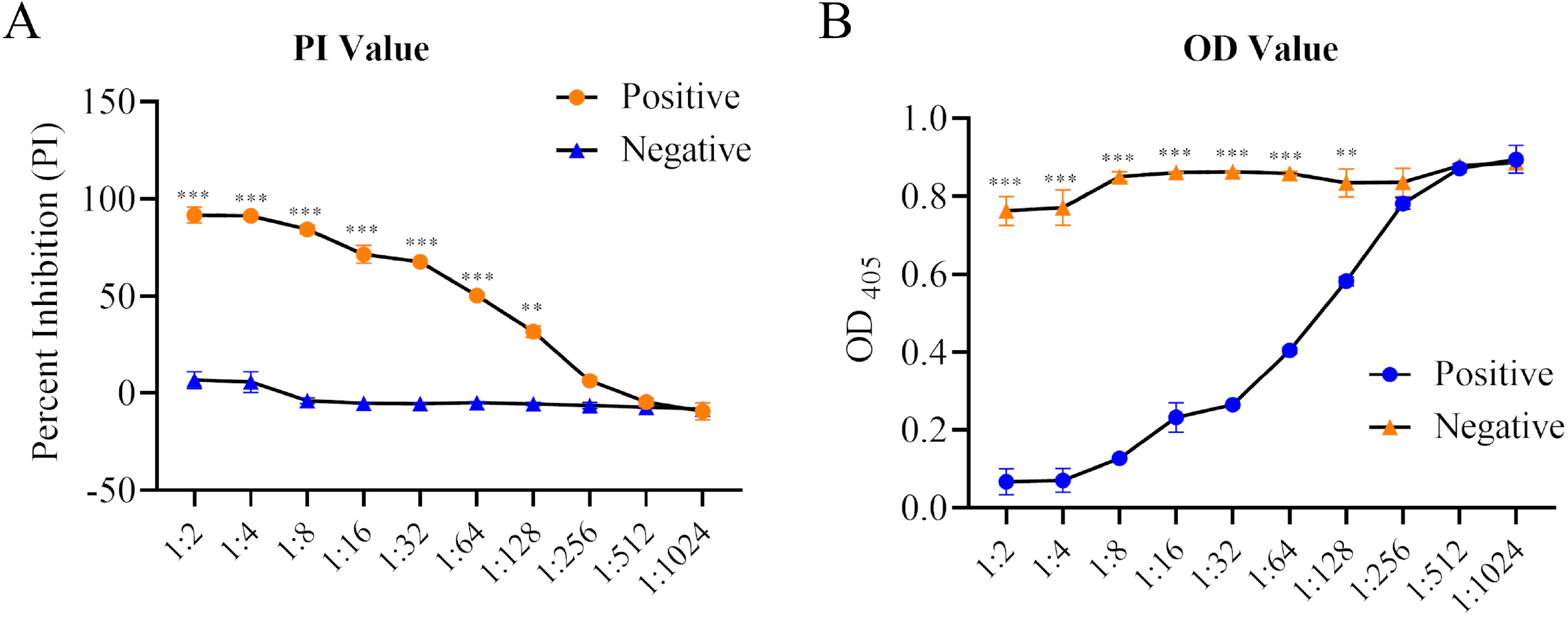
Analytical sensitivity of bELISA. Two-fold serial dilutions of the high-positive and negative cat serum control standards were run in parallel. Each dilution was tested in duplicates. OD values (**A**) or percentage of inhibition (PI) values (**B**) were calculated and displayed in Y-axis. Differences under each dilution were analyzed by one-way analysis of variance (ANOVA) using GraphPad Prism 6 software (GraphPad, La Jolla, CA). P-values were indicated by asterisks. ** P < 0.01, *** P < 0.001.

#### Diagnostic sensitivity and specificity of bELISA

To evaluate the diagnostic sensitivity and specificity of the mAb-based bELISA, a panel of serum samples with known antibody status was tested, including 45 positives and 88 negatives collected from cat, ferret, mink, and deer. Before testing in bELISA, all serum samples were analyzed by serum neutralization assay to confirm the antibody status. The bELISA result showed that a cut-off PI value of 17.60% produced a maximized diagnostic sensitivity of 97.8% (95% confidence interval: 88.2-99.9%) and diagnostic specificity of 98.9% (95% confidence interval: 93.8-100%) (Figure 4A). Subsequently, a single-graph ROC analysis was conducted by comparing false-positives (1 - diagnostic specificity) and true-positives (diagnostic sensitivity). The area under the curve (AUC) represents the overall accuracy of the test. An AUC of 1 indicates a perfect test, and above 0.9 indicates high accuracy. The AUC of #127-3 mAb-based bELISA was 0.998 (p < 0.001) with a 95% confidence interval of 97%–100%, demonstrating the high accuracy of the assay (Figure 4B).

**Figure 4.**
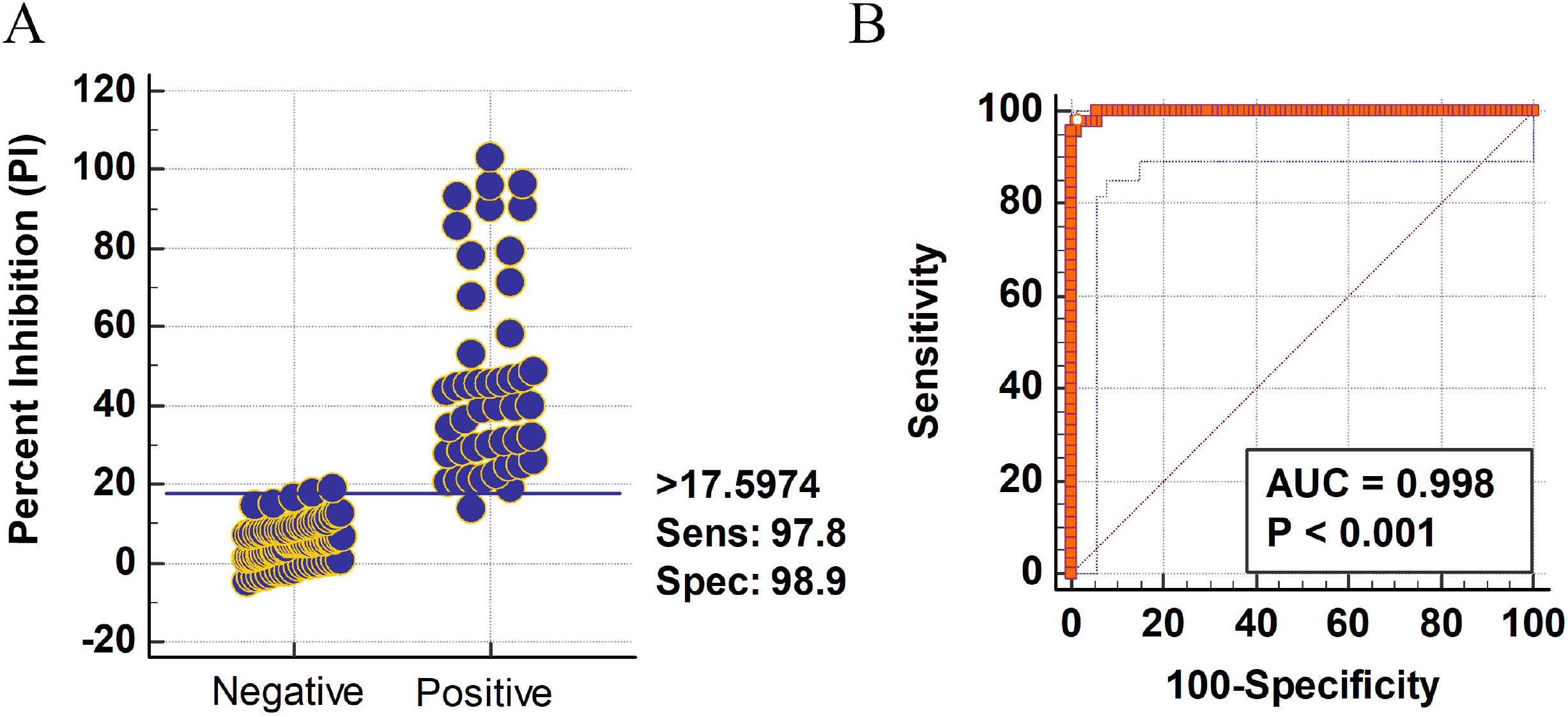
Determination of diagnostic sensitivity and specificity. Receiver operating characteristic (ROC) analysis **(A)** and the interactive plot of diagnostic sensitivity and specificity **(B)** were calculated using 45 known-positive serum samples and 88 known-negative serum samples collected from different animal species, including cat, ferret, mink, and deer. A horizontal line between the positive and negative populations in panel A represents the cutoff value that produces the optimal diagnostic sensitivity and specificity. ROC analysis was conducted by using MedCalc software (version 10.4.0.0, MedCalc Software, Mariarke, Belgium).

#### Repeatability of bELISA

Repeatability determines the ability of an assay to produce similar results from multiple preparations and runs of a same sample. In this study, repeatability of #127-3 mAb-based bELISA was assessed by running a single lot of medium-positive control serum standard. The percentage of coefficient of variation (% CV) was calculated to measure the repeatability. The results showed that within plate % CV was 5.15% (mean value of 55.37% + standard deviation of 2.84%), between-plate % CV within one run was 6.95% (mean value of 55.37% + standard deviation of 3.85), while the between runs % CV was 7.23% (mean value of 55.37% + standard deviation of 4%). The values of % CV below 10% indicate that the #127-3 mAb-based bELISA is highly repeatable (18, 21).

### Detection of seroconversions in SARS-CoV-2 infected cats

Next, we applied the bELISA to investigate the dynamics of anti-N antibody response in SARS-CoV-2 infected cats. Serum samples were collected from our previous study (19), in which 3 groups of cats (n = 8) were experimentally inoculated with each of the SARS-CoV-2 variants (B.1, Delta, Omicron). Serum samples were collected at 0, 3, 5, 7 and 14 days post infection. This set of samples was tested by bELISA and results showed that anti-N antibody response was detected as early as 7 dpi for B.1 and Delta variants, then dramatically increased to a high level (PI = 47.03% for B.1, PI = 71.42% for Delta variant) at 14 dpi (Figure 5A). Omicron variant-induced antibody response (PI = 27.87%) was detected at a late time point (14 dpi). Overall, Delta variant induced the highest antibody response compared to B.1 and Omicron variants. This result is consistent with that of virus neutralization assay. The same trend of dynamics was also observed for serum neutralizing acitivities against the live virus of Delta variant (B.1.617.2) using the same set of serum samples (Figure 5B).

**Figure 5.**
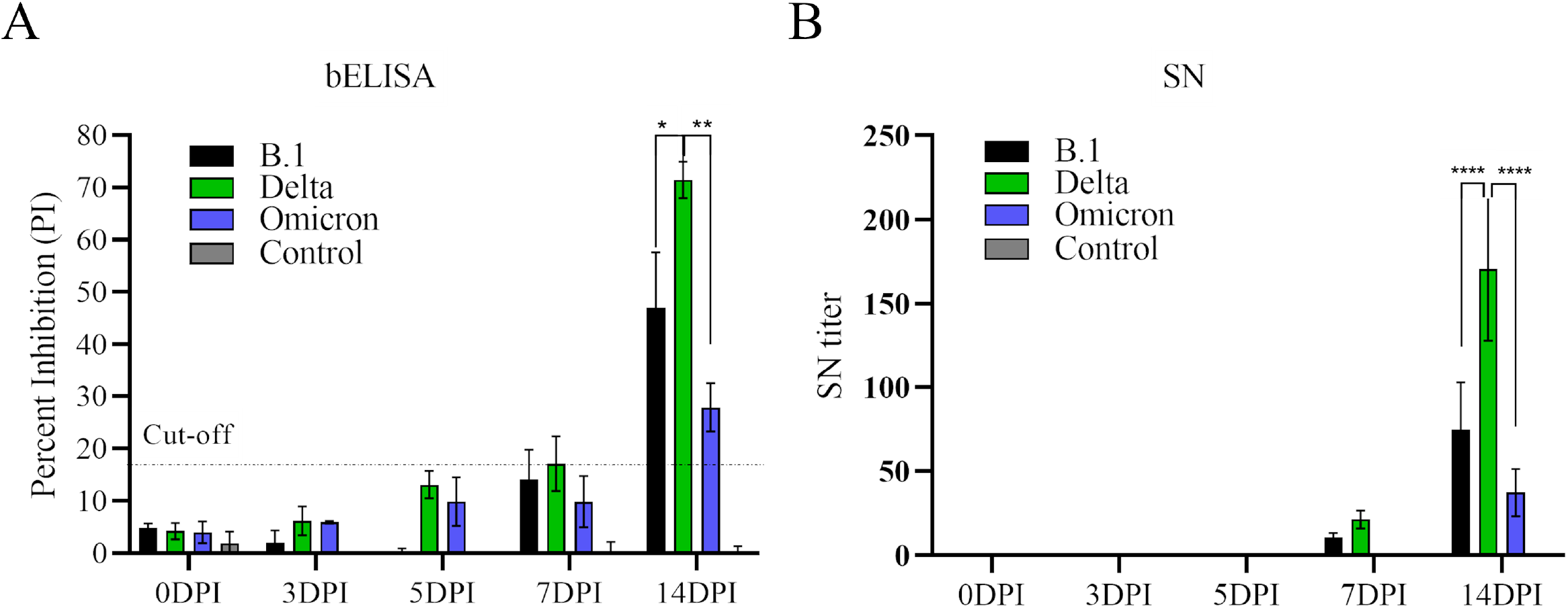
Dynamics of antibody response in cats infected by different SARS-CoV-2 variants. A total of 24 domestic cats were divided into four groups, in which each group was inoculated with one of the SARS-CoV-2 variants (B.1, Delta, Omicron) and group four was mock-inoculated with cell culture medium. Serum samples were collected before infection and 3, 5, 7, 17 days post infection (DPI). (**A**) bELISA test to measure the antibody response through the time course study. The dashed line represents the cutoff value (17.60%) of the assay. (**B**) Serum neutralization assay. The assay was performed using SARS-CoV-2 Delta variant (B.1.617.2). Neutralizing antibody titer was calculated as the reciprocal of the highest serum dilution that generated 100% neutralization of SARS-CoV-2 infection. Statistical differences between each group within each time point were calculated using one-way analysis of variance (ANOVA). * P < 0.05, ** P < 0.01.

### Application of bELISA in pet animals with clinical diseases

We further applied the bELISA for detection of SARS-CoV-2 infection in pet animals. Serum samples were collected from three dogs in a pet clinic. These dogs were experiencing clinical signs of respiratory diseases. The bELISA result showed that two dogs (Dog-1 and Dog-2) were positive for SARS-CoV-2 antibodies with PI values of 18.66% and 46.33% respectively, while the third dog was negative for specific anti-N antibody with a PI value of 2.54% (Figure 6A). The result was further confirmed by serum neutralizing test at USDA NVSL laboratory. The result showed neutralizing titers of 92.86%, 37.04%, and 5.69% for dog 1, 2, 3, respectively (Figure 6A). Dog-2 exhibited long-term illness, and returned back to the pet clinic periodically. Serum samples were collected from this dog during each examination in the clinic from February to August, 2022. The bELISA detected the increased antibody titer in 15 days (February 22^nd^, 2022; PI = 77.54%) after the first examination (February 7^th^, 2022; PI = 46.33%). The titer was decreased at the third examination (March 10^th^, 2022; PI = 48.45%). At the fourth examination (August 2^nd^, 2022), 176 days from the first examination, lower level of antibody titer (PI = 31.15%) was still detected (Figure 6B).

**Figure 6.**
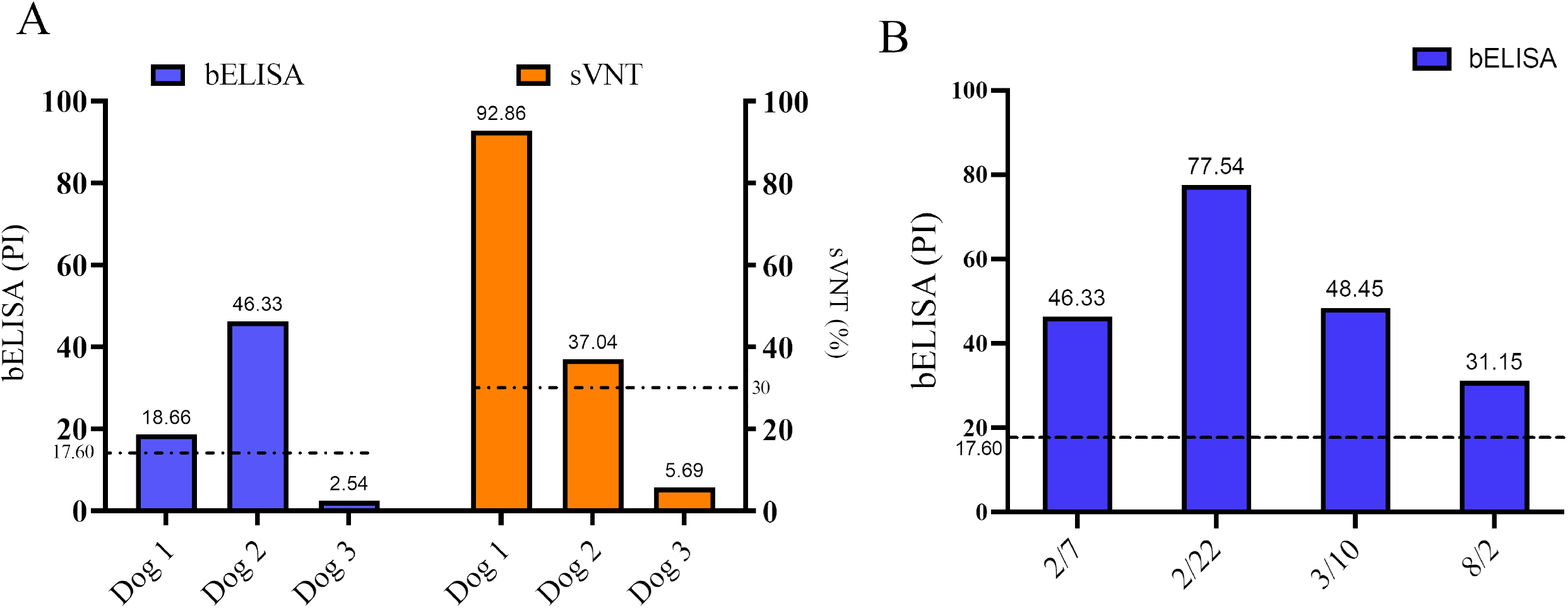
Detection of SARS-CoV-2 specific antibody response in dogs with clinical diseases. **(A)** Serum samples from three dogs were tested by bELISA and sVNT. bELISA results were presented in blue bar, while sVNT results were presented in orange bar. X-axis shows individual dogs and values for both assays were presented on top of column. (**B**) Serum antibody titers in Dog 2 tested by bELISA through a time course study. Dashed lines represent the cut-off values for bELISA (17.60%) and sVNT (30%).

## DISCUSSION

The COVID-19 pandemic has emphasized the critical role of effective diagnostics in the response to outbreaks. Diagnostic tools for active surveillance and monitoring of SARS-CoV-2 are essential for the successful control of the pandemic. Reverse zoonotic transmission of this virus from animals to humans has also been reported, which highlights the need for accurate diagnostic tools to be used at the human-animal interface (22–24). Sera-based diagnostics applicable for large-scale field surveillance in all animal species becomes important to understand mechanism of zoonotic transmission. MAbs are key reagents for detection of viral infections and study the viral pathogenesis. Therefore, the goal of this study was to produce a panel of mAbs against SARS-CoV-2 N protein and develop a mAb-based bELISA for sera-surveillance in an animal species-independent manner.

Utilizing hybridoma technology, a panel of mAbs recognizing different epitopes of SARS-CoV-2 N protein was generated. It allows us to select the suitable mAb for bELISA development. The mAb #127-3 was characterized to have strong reactivity in cells infected by different SARS-CoV-2 variants. It did not cross-react with the N protein of other human/animal coronaviruses that we tested, which contributes to the high specificity of the mAb-based bELISA developed thereafter. Due to the unique design of bELISA, high specificity is expected as reported previously for assays targeting African swine fever virus (21) and porcine reproductive and respiratory syndrome virus (18). Our mAb-based bELISA achieved high sensitivity (97.8%) and specificity (98.9%), which is comparable to the current commercially available serological tests. The Abbott assay (SARS-CoV-2 IgG assay, Abbott, Chicago, IL, US) was reported to reach 92.7% sensitivity and 99.9% specificity, the DiaSorin assay (LIAISON SARS-CoV-2 S1/S2 IgG, DiaSorin, Saluggia, Italy) has 96.2% sensitivity and 98.9% specificity, and the Roche assay (Elecsys Anti-SARS-CoV-2 assay, Roche, Basel, Switzerland) has 97.2% sensitivity and 99.8% specificity in human serum samples (25). Similarly, the SARS-CoV-2 surrogate neutralization test achieves 96% sensitivity and 99.93% specificity (26). Much higher sensitivity can be achieved in symptomatic individuals and those in the late phase of infection due to robust production of antibody responses (27).

Current available serological assays for SARS-CoV-2, include ELISAs, are targeting host antibody response against N or S protein, and most of them are specifically designed for human samples. For example, the Abbott and Roche assays target N protein, while the DiaSorin assay targets S protein. They all primarily are designed for testing human samples and require species-specific secondary antibodies for testing the samples from a specific animal species (25). Notably, the surrogate neutralization test adapted the ELISA format to block bindings between coating ACE2 receptor and HRP conjugated Spike/RBD proteins, which is a cell- and virus-free assay and capable of screening serum samples from all host species (16). However, measuring neutralizing antibodies has to accommodate different variants, since frequent mutations in S protein leads to potential mis-binding of ACE2 and S protein. The mAb-based bELISA developed in this study targets N protein, which is highly conserved across different variants of SARS-CoV-2, thus has less probability to be affected by emerging variants. In addition, due to the abundant presence of N protein, immunoassays targeting N protein are more sensitive than that targeting S protein, especially during the early infection stage (28–31). Previous studies showed that serum SARS-CoV-2 N protein could be a diagnostic marker for detection of early infections (32–34). In the case of SARS coronavirus, N protein could be detected in serum samples from 95% SARS patients at just 3 days after symptom onset (35). Consistently, our bELISA was able to detect antibodies against B.1 and Delta variants in cats at 7 days post infection. Furthermore, in combination with an S protein-based test, the N protein-based bELISA is capable of differentiating between infected and vaccinated animals when an S protein-based COVID-19 vaccine is used.

We further applied the bELISA to diagnose pet animals with clinical illness. Two dogs were tested positive on the bELISA and results were further confirmed by virus neutralizing assays. Oropharyngeal samples were further collected from both dogs and quantitative RT-PCR was conducted by the USDA NVSL laboratory. The result showed that CT value of Dog-1 was 37.66 with N1 primer set and negative with N2 primer set, while CT value of Dog-2 was 31.32 with N1 primer set and 33.99 with N2 primer set for SARS-CoV-2 nucleic acid detection. These results fall into “suspect” category according to CDC guidelines (36). The owner of Dog-1 was diagnosed as COVID-19 positive in January 2022, suggesting that the dog might have been exposed to the SARS-CoV-2 from the owner and subsequently developed the specific antibody response. The samples that we tested were collected in early February 2022, which might have been about 2-3 weeks after potential exposure to the virus. At this stage, the animal should have already passed the peak time for shedding the virus and developed specific immune response against the viral infection (37). This could explain our observation of a “suspect” level of nucleic acid detected in RT-PCR test, but high level of antibody detected in bELISA and virus neutralizing test. Interestingly, antibody response in Dog-2 lasted for about 6 months. This result is consistent with previous findings in humans, in which a longitudinal analysis of antibody dynamics in COVID-19 convalescents demonstrated that both neutralizing and non-neutralizing antibodies can still be detected over 8 months post-symptom onset, although the titer was substantially decreased (38–40).

In summary, the panel of mAbs generated in this study provides valuable reagents for disease diagnostics and viral pathogenesis studies. The mAb-based bELISA could be a useful tool for field surveillance to determine the prevalence of COVID-19 in animal populations and identify potential new animal reservoirs.

## MATERIALS AND METHODS

### Cells, viruses, and viral genes

Vero-E6 and MARC-145 cells were maintained in minimum essential medium (Thermo Fisher Scientific, Waltham, MA) supplemented with 10% heat-inactivated fetal bovine serum (Sigma-Aldrich, Burlington, MA) and antibiotics (100 μg/mL streptomycin, 100 U/mL penicillin, and 0.25 μg/mL fungizone) at 37°C with 5% CO_2_.

The SARS-CoV-2 isolates used in this study were obtained from residual de-identified human anterior nares or nasopharyngeal secretions (Institutional review board [IRB] at Cayuga Health System [protocol 0420EP] and Cornell University [protocol 2101010049]). The SARS-CoV-2 D614G (B.1 lineage) New York-Ithaca 67-20 (NYI67-20), Alpha (B.1.1.7) New York City 853-21 (NYC853-21), and Delta (B.1.617.2 lineage) NYI31-21 isolates, were propagated in Vero E6/TMPRSS2 cells, whereas the Omicron BA.1.1 (B.1.1.529) NYI45-21 isolate was propagated in Vero E6 cells in BSL3 laboratory conditions at the Animal Health Diagnostic Center (AHDC) Research Suite at Cornell University. The SARS-CoV-2 full-length N gene of Wuhan-hu-1 isolate (GenBank # NC 045512.2) was synthesized (GenScript, Piscataway, NJ) and cloned in the pET-28a (+) vector (Novagen, Madison, WI) or pCAGGS vector (provided by Dr. Adolfo Garcia-Sastre at the Icahn School of Medicine at Mount Sinai in New York City) (41). In addition, N genes of common coronaviruses that infect SARS-CoV-2 susceptible animal hosts were synthesized. Each synthetic gene was fused with a Flag tag (DYKDDDDK) at its C terminus and cloned into a plasmid vector pTwist-CMV-BetaGlobin (Twist Bioscience, San Francisco, CA). The synthesized genes were derived from human coronavirus OC43 (HCoV-OC43; GenBank ID, AY585228.1), human coronavirus NL63 (HCoV-NL63; GenBank ID, AY567487.2), human coronavirus 229E (HCoV-229E; GenBank ID, NC_002645.1), human coronavirus HKU1 (HCoV-HKU1; GenBank ID, NC_006577.2), severe acute respiratory syndrome coronavirus (SARS-CoV; GenBank ID, AY278741.1), middle east respiratory syndrome coronavirus (MERS-CoV; GenBank ID, NC_019843.3), feline infectious peritonitis virus (FIPV; GenBank ID, AY994055.1), feline coronavirus (FCoV; GenBank ID, EU186072.1), canine coronavirus type I (CCoV-type I; GenBank ID, KP849472.1), canine coronavirus type II (CCoV-type II; GenBank ID, KC175340.1), ferret systemic coronavirus (FRSCV; GenBank ID, GU338456.1), and mink coronavirus (MCoV; GenBank ID, HM245925.1).

### Recombinant protein preparation

Recombinant N protein of SARS-CoV-2 was expressed in BL21 *E.coli* as a polyhistidine (6x His-tagged) fused protein. The antigen was produced and purified by following a method described in our previous study (18). Purified proteins were dialyzed using 1x phosphate-buffered saline (PBS) solution under 4°C for three times and then concentrated by polyethylene glycol 8000 (Thermo Fisher Scientific, Waltham, MA).

### Monoclonal antibody (mAb) production

BALB/c mice were immunized with recombinant N protein at a dose of 50-100 μg per mouse and further boosted 2-3 times at an interval of two to three weeks. At three days after the final boost, mice splenocytes were collected and fused with NS-1 myeloma cells to generate hybridoma cells. Specific anti-N antibody-secreting hybridomas were screened by using immunofluorescent assays (see below). Selected hybridomas were expanded in large tissue culture flask. Cell culture supernatants containing specific anti-N mAb were harvested and concentrated using Pierce™ Saturated Ammonium Sulfate Solution (Thermo Fisher Scientific, Waltham, MA). Biotinylation of the mAb was performed using a Biotin Conjugation Kit by following the manufacturer’s instruction (Abcam, Cambridge, MA). The SARS-CoV-2 N-specific mAb B6G11 was previously developed in Diel lab (20).

### Immunofluorescent assay (IFA)

For screening hybridomas and performing antibody cross-reactivity test with other coronaviruses, MARC-145 cells were seeded in 96-well cell culture plates and transfected with plasmid DNA expressing N protein of the corresponding coronavirus. Transfection was performed using TransIT®-LT1 Transfection Reagent (Mirus Bio, Madison, WI). At 48 hours post transfection, cells were fixed with 80% acetone (Thermo Fisher Scientific, Waltham, MA) for 10 min at room temperature. Cell monolayers were incubated with the primary mAb at 37°C for 1 hour, followed by incubation with the secondary antibody, Alexa Fluor 488 AffiniPure goat anti-mouse IgG (H+L) (Jackson Immuno Research, West Grove, PA). Immunofluorescent signals were visualized with an inverted immunofluorescent microscope (LMI6000, LAXCO, Mill Creek, WA). To confirm the reactivity and specificity of the anti-N mAb, Vero E6 Cells were infected with different SARS-CoV-2 variants. At 24 hours post infection, cells were fixed with 3.7% formaldehyde solution in PBS for 30 min followed by permeabilization with 0.1% Triton-X-100 in PBS for 10 min at room temperature. After 3 consecutive washing steps with PBS, anti-SARS-CoV-2 mAbs diluted in blocking solution (1% BSA in PBS) were added to the cells and incubated for 1 hour at 37 °C in a humidified chamber. Cells were washed again and incubated under the same conditions with goat anti-mouse IgG AlexaFluor 594. Cell nuclei were stained with DAPI and image acquisition was performed with an inverted immunofluorescent microscope.

### Western blot

MARC-145 cells were transfected with plasmid DNA of pCAGGS-N that contains SARS-CoV-2 full-length N gene. At 48 hours post transfection, cells were harvested with Pierce™ IP Lysis Buffer (Thermo Fisher Scientific, Waltham, MA) containing Protease Inhibitor Cocktail (Sigma-Aldrich, St. Louis, MO). Western blot analysis was performed using the method as we described previously (21). The membrane was probed with specific anti-N mAb as the primary antibody and detected by IRDye 800CW goat anti-mouse IgG (H+L) (Li-Cor Biosciences, Lincoln, NE) as the secondary antibody. Protein blots were imaged using an Odyssey Fc imaging system (Li-Cor Biosciences, Lincoln, NE).

### Immunoprecipitation

MARC-145 cells transfected with the recombinant pCAGGS-N plasmid were lysed in Pierce™ IP Lysis Buffer (Thermo Fisher Scientific, Waltham, MA), and then mixed with each of the purified anti-N mAbs. Immune-complexes were precipitated by Protein A/G magnetic beads (Thermo Fisher Scientific, Waltham, MA). Precipitated proteins were separated by SDS-PAGE and analyzed by Western blot as described previously (42).

### Serum neutralization test

Serum neutralization (SN) assay was performed under BSL-3 laboratory conditions at Cornell University. Two-fold serial dilutions (1:8 to 1:1024) of cat serum samples were incubated with SARS-CoV-2 Delta variant (B.1.617.2) (100-200 TCID50/well) for 1 hour at 37°C. Following incubation of serum and virus, 50 μL of a cell suspension of Vero E6 cells was added to each well of a 96-well plate and incubated for 48 hours at 37°C with 5% CO_2_. Cells were fixed and subjected to IFA as previously described previously (19). The neutralizing antibody titer was calculated as the reciprocal of the highest serum dilution that generated 100% neutralization of SARS-CoV-2 infection. Samples with antibody titer less than 1:8 were considered as negative.

The surrogate virus neutralization test (sVNT) was performed at USDA National Veterinary Services Laboratory (NVSL) at Ames, Iowa. A cPass™ SARS-CoV-2 Neutralization Antibody Detection Kit (GenScript, Piscataway, NJ) was used and the test was performed following the instructions of the manufacture. Briefly, 10 μL of serum sample was diluted with 90 μL of sample dilution buffer, followed by taking 60 μL of diluted sample to react with 60 μL HRP-conjugated RBD solution. The mixture of sample and HRP-RBD was incubated at 37°C for 30 minutes. The incubated mixture (100 uL) was added to the plate wells that were pre-coated with hACE2 antigen and then incubate at 37°C for 15 minutes. Wells were washed for three times, followed by addition of 100 μL TMB Solution to each well and incubation in dark at room temperature for 15 minutes. Finally, 50 μL of Stop Solution was added to each well and plate was read at 450 nm using a spectrophotometer. The percent signal inhibition for detecting neutralizing antibodies were calculated and the sample was determined as neutralizing antibody positive if the percent signal inhibition was more than 30%.

### Sample sources

The control serum standards used for ELISAs were created using serum samples collected from our previous cat experiment (19). The positive control serum was collected from cats that were experimentally inoculated with SARS-CoV-2 D614G (B.1), Delta (B.1.617.2), or Omicron (B.1.1.529) variant at 14 days post infection (dpi), while the negative control serum was collected from negative control cats. Large quantities of positive sera were pooled into a single lot of positive control serum, and large quantities of the negative sera were pooled into a single lot of negative control serum. The high-, medium-, and low-positive control serum standards were created by spiking the positive control serum into the negative control serum to generate the desired antibody titers in the ELISAs.

For assay validation, four sets of animal serum samples with known infection status were used. The first set contained 17 positive and 43 negative serum samples collected from cats infected with SARS-CoV-2 D614G (B.1), Delta (B.1.617.2), or Omicron (B.1.1.529) strain in study described previously (19). The second set contained 10 positive and 37 negative serum samples collected from SARS-CoV-2 isolate NYI67-20 (B.1 lineage) infected ferrets (43). The third set contained 5 positive and 8 negative serum samples collected from SARS-CoV-2 (lineage B) infected deer (44). The fourth set included 13 positive mink serum samples. The antibody status of all the serum samples used for bELISA validation was confirmed by serum neutralizing assay as described above.

The capability of the bELISA to detect the seroconversion was evaluated using samples collected from a cat experiment that we reported previously (19). Serum samples were collected at 0, 3, 5, 7, 14 days post infection (dpi).

To apply the bELISA in the diagnosis of clinical animals, serum and oropharyngeal samples were collected from 3 dogs at a pet clinic in Illinois. Dog-1 was a 6-year-old, male neutered, Samoyed. At the time (Feb 7, 2022) that samples were collected for SARS-CoV-2 tests, the dog had clinical signs of coughing and sneezing for about three weeks and was tested positive for *Mycoplasma*. Dog-2 was a 5.5-month-old, male, Great Dane mix. The dog started showing clinical signs of coughing, vomiting, decreased appetite, and extreme lethargy in late January of 2022. Samples from Dog-2 were collected on February 7^th^ for testing. Dog-3 was 14-year-old, female sprayed, mixed breed dog, displaying coughing and sneezing on March 3, 2022. She also had a history of airway disease. Samples were collected on March 10, 2022.

### Procedure for blocking ELISA and indirect ELISA

Both ELISAs were performed using our previously described methods with modifications (21, 45). The bELISA could detect antibodies from multiple animal species by allowing sample antibody binding to the coated antigen on the ELISA plate first, followed by adding biotin-conjugated mAb. If presence of anti-N antibodies in the animal serum, they will bind to the N antigen and block the binding of biotinylated anti-N mAb to the N antigen. The mAb will be washed away and no color signal will be developed in the subsequent steps. If there is no anti-N antibodies present in the animal serum, the biotinylated anti-N mAb will bind to the N antigen, then the HRP-conjugated streptavidin will be added and bind to the biotin that conjugated to mAb. HRP substrate will be added to develop the color signal. Thus, the amount of anti-N antibodies in the testing sample is inversely proportional to the level of color signal. To conduct the bELISA test, initially, the odd number columns in Immulon 2HB plate (Thermo Fisher Scientific, Waltham, MA, USA) were coated with recombinant N protein (175ng) diluted in antigen coating buffer (ACB; 35 mM sodium bicarbonate and 15 mM sodium carbonate, PH 8.8). The even number columns in the plate were added with ACB only as the background control. The plate was incubated at 37°C for 1 hour and then 4°C overnight. After blocking with 2% bovine serum albumin (BSA; Thermo Fisher Scientific, Waltham, MA, USA) in PBST (0.05% Tween 20 in 1x phosphate-buffered saline) at 37°C for 1 hour, the plate was washed three times by PBST using the automated microplate washer (BioTek, Winooski, VT). The test serum samples were diluted 1:4 with 2% BSA and added into both coated and uncoated wells. The internal control standards (100 ul; high-, medium-, low-positive, and negative) were added in duplicates. After incubation for 1 hour at 37°C, 100 μL of biotinylated mAb (clone #127-3) was added and incubated at 37°C for another 30 min. The plate was washed for three times and incubated with 100 uL of streptavidin poly-HRP (1:2000 dilution; Thermo Fisher Scientific, Waltham, MA) at room temperature for 1 hour. After wash with PBST, 100 μL of ABTS peroxidase substrate (KPL, Gaithersburg, MD) was added for color development. The colorimetric reaction was stopped by equal volume of ABTS stop solution (KPL, Gaithersburg, MA) in 5 min and color intensity was quantified at 405 nm using a SpectraMax® iD5 microplate reader (Molecular Devices, San Jose, CA). The percentage of inhibition was calculated using the following formula:

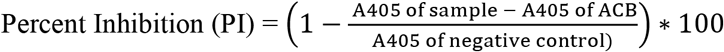

For indirect ELISA, the plate was coated using the same method as that of bELISA. After blocking with 5% non-fat milk in PBST, serum samples (1:400 dilution in 5% non-fat milk) and internal control standards were loaded on the plate and incubated at 37°C for 1 hour. The plate was washed for three times and then added 100 uL of HRP-conjugated goat anti-feline IgG (H+L) secondary antibody (1:5000 dilution; Thermo Fisher Scientific, Waltham, MA) for incubation another hour at 37°C. After washing of the plate, colorimetric reaction was developed by adding ABTS peroxidase substrate and stopped by ABTS stop solution. Color development was quantified using the SpectraMax® iD5 microplate reader (Molecular Devices, San Jose, CA).

### Validation of N protein-based blocking ELISA

For analytical sensitivity analysis of the bELISA, two-fold serial dilutions of the high-positive and negative serum standards were tested in triplicate and differences between different dilutions of the control serum were evaluated by one-way analysis of variance (ANOVA) using Prism software version 6 (GraphPad Software, San Diego, California). A *p*-value of less than 0.01 (**) was considered as statistically significant.

To determine the optimal diagnostic sensitivity and specificity, the four sets of known-status animals serum samples mentioned above were subject to bELISA test. Calculations of the assay performance were conducted using MedCalc®, version 10.4.0.0 (MedCalc® Software, Mariarke, Belgium). The cutoff of bELISA was defined as the PI value that was able to produce the maximized diagnostic sensitivity and specificity. In addition, Receiver operating characteristic (ROC) analysis was performed using the same software to assess the overall accuracy of the assay.

The assay repeatability was determined by running repeated samples of the medium-positive control. Assay precisions were calculated as 40 replicates in one plate for within-plate level, 3 plates in one run for between-plate level, and 3 consecutive runs for between-run level. Means, standard deviations, and percent coefficient of variation (%CV) values were calculated using Control Chart Pro Plus software (ChemSW, Inc., Fairfield Bay, AR, USA).

## ACKNOWLEDGMENTS

We thank Dr. Mia K. Torchetti from USDA National Veterinary Services Laboratories for testing the dog samples. This project was supported by the National Institute of Health (Grant# R01AI166791). Fangfeng Yuan was partially supported by Illinois Distinguished Fellowship for graduate student at the University of Illinois.

## Notes

### Competing Interest Statement

The authors have declared no competing interest.

## REFERENCES

1. Rambaut A, Holmes EC, O’Toole Á, Hill V, McCrone JT, Ruis C, du Plessis L, Pybus OG. 2020. A dynamic nomenclature proposal for SARS-CoV-2 lineages to assist genomic epidemiology. Nature Microbiology 5:1403–1407.

2. Enjuanes L, Gorbalenya AE, de Groot RJ, Cowley JA, Ziebuhr J, Snijder EJ. 2008. Nidovirales. Encyclopedia of Virology doi:10.1016/B978-012374410-4.00775-5:419-430.

3. V’kovski P, Kratzel A, Steiner S, Stalder H, Thiel V. 2021. Coronavirus biology and replication: implications for SARS-CoV-2. Nature Reviews Microbiology 19:155–170.

4. Heffron AS, McIlwain SJ, Amjadi MF, Baker DA, Khullar S, Armbrust T, Halfmann PJ, Kawaoka Y, Sethi AK, Palmenberg AC, Shelef MA, O’Connor DH, Ong IM. 2021. The landscape of antibody binding in SARS-CoV-2 infection. PLOS Biology 19:e3001265.

5. Damas J, Hughes GM, Keough KC, Painter CA, Persky NS, Corbo M, Hiller M, Koepfli K-P, Pfenning AR, Zhao H, Genereux DP, Swofford R, Pollard KS, Ryder OA, Nweeia MT, Lindblad-Toh K, Teeling EC, Karlsson EK, Lewin HA. 2020. Broad host range of SARS-CoV-2 predicted by comparative and structural analysis of ACE2 in vertebrates. Proceedings of the National Academy of Sciences 117:22311.

6. Sailleau C, Dumarest M, Vanhomwegen J, Delaplace M, Caro V, Kwasiborski A, Hourdel V, Chevaillier P, Barbarino A, Comtet L, Pourquier P, Klonjkowski B,Manuguerra J-C, Zientara S, Le Poder S. 2020. First detection and genome sequencing of SARS-CoV-2 in an infected cat in France. Transboundary and emerging diseases 67:2324–2328.

7. McAloose D, Laverack M, Wang L, Killian Mary L, Caserta Leonardo C, Yuan F, Mitchell Patrick K, Queen K, Mauldin Matthew R, Cronk Brittany D, Bartlett Susan L, Sykes John M, Zec S, Stokol T, Ingerman K, Delaney Martha A, Fredrickson R, Ivančić M, Jenkins-Moore M, Mozingo K, Franzen K, Bergeson Nichole H, Goodman L, Wang H, Fang Y, Olmstead C, McCann C, Thomas P, Goodrich E, Elvinger F, Smith David C, Tong S, Slavinski S, Calle Paul P, Terio K, Torchetti Mia K, Diel Diego G, Meng X-J. From People to Panthera: Natural SARS-CoV-2 Infection in Tigers and Lions at the Bronx Zoo. mBio 11:e02220–20.

8. Sharun K, Tiwari R, Saied AA, Dhama K. 2021. SARS-CoV-2 vaccine for domestic and captive animals: An effort to counter COVID-19 pandemic at the human-animal interface. Vaccine 39:7119–7122.

9. Chandler JC, Bevins SN, Ellis JW, Linder TJ, Tell RM, Jenkins-Moore M, Root JJ, Lenoch JB, Robbe-Austerman S, DeLiberto TJ, Gidlewski T, Kim Torchetti M, Shriner SA. 2021. SARS-CoV-2 exposure in wild white-tailed deer (*Odocoileus virginianus*). Proceedings of the National Academy of Sciences 118:e2114828118.

10. Oreshkova N, Molenaar RJ, Vreman S, Harders F, Oude Munnink BB, Hakze-van der Honing RW, Gerhards N, Tolsma P, Bouwstra R, Sikkema RS, Tacken MG, de Rooij MM, Weesendorp E, Engelsma MY, Bruschke CJ, Smit LA, Koopmans M, van der Poel WH, Stegeman A. 2020. SARS-CoV-2 infection in farmed minks, the Netherlands, April and May 2020. Euro surveillance: bulletin Europeen sur les maladies transmissibles = European communicable disease bulletin 25:2001005.

11. Medkour H, Catheland S, Boucraut-Baralon C, Laidoudi Y, Sereme Y, Pingret J-L, Million M, Houhamdi L, Levasseur A, Cabassu J, Davoust B. 2021. First evidence of human-to-dog transmission of SARS-CoV-2 B.1.160 variant in France. Transboundary and emerging diseases doi:10.1111/tbed.14359:10.1111/tbed.14359.

12. Barroso-Arévalo S, Rivera B, Domínguez L, Sánchez-Vizcaíno JM. 2021. First Detection of SARS-CoV-2 B.1.1.7 Variant of Concern in an Asymptomatic Dog in Spain. Viruses 13:1379.

13. Kevadiya BD, Machhi J, Herskovitz J, Oleynikov MD, Blomberg WR, Bajwa N, Soni D, Das S, Hasan M, Patel M, Senan AM, Gorantla S, McMillan J, Edagwa B, Eisenberg R, Gurumurthy CB, Reid SPM, Punyadeera C, Chang L, Gendelman HE. 2021. Diagnostics for SARS-CoV-2 infections. Nature materials 20:593–605.

14. Deeks JJ, Dinnes J, Takwoingi Y, Davenport C, Spijker R, Taylor-Phillips S, Adriano A, Beese S, Dretzke J, Ferrante di Ruffano L, Harris IM, Price MJ, Dittrich S, Emperador D, Hooft L, Leeflang MM, Van den Bruel A, Cochrane C-DTAG. 2020. Antibody tests for identification of current and past infection with SARS-CoV-2. The Cochrane database of systematic reviews 6:CD013652-CD013652.

15. Liu K-T, Han Y-J, Wu G-H, Huang K-YA, Huang P-N. 2022. Overview of Neutralization Assays and International Standard for Detecting SARS-CoV-2 Neutralizing Antibody. Viruses 14:1560.

16. Fenwick C, Turelli P, Pellaton C, Farina A, Campos J, Raclot C, Pojer F, Cagno V, Nusslé SG, D’Acremont V, Fehr J, Puhan M, Pantaleo G, Trono D. 2021. A high-throughput cell-and virus-free assay shows reduced neutralization of SARS-CoV-2 variants by COVID-19 convalescent plasma. Science Translational Medicine 13:eabi8452.

17. Henriques AM, Fagulha T, Barros SC, Ramos F, Duarte M, Luís T, Fevereiro M. 2016. Development and validation of a blocking ELISA test for the detection of avian influenza antibodies in poultry species. Journal of Virological Methods 236:47–53.

18. Ferrin NH, Fang Y, Johnson CR, Murtaugh MP, Polson DD, Torremorell M, Gramer ML, Nelson EA. 2004. Validation of a blocking enzyme-linked immunosorbent assay for detection of antibodies against porcine reproductive and respiratory syndrome virus. Clinical and diagnostic laboratory immunology 11:503–514.

19. Martins M, do Nascimento Gabriela M, Nooruzzaman M, Yuan F, Chen C, Caserta Leonardo C, Miller Andrew D, Whittaker Gary R, Fang Y, Diel Diego G. 2022. The Omicron Variant BA.1.1 Presents a Lower Pathogenicity than B.1 D614G and Delta Variants in a Feline Model of SARS-CoV-2 Infection. Journal of Virology 96:e00961–22.

20. Carvallo FR, Martins M, Joshi LR, Caserta LC, Mitchell PK, Cecere T, Hancock S, Goodrich EL, Murphy J, Diel DG. 2021. Severe SARS-CoV-2 Infection in a Cat with Hypertrophic Cardiomyopathy. Viruses. 13(8):doi:10.3390/v13081510.

21. Yuan F, Petrovan V, Gimenez-Lirola LG, Zimmerman JJ, Rowland RRR, Fang Y. 2021. Development of a Blocking Enzyme-Linked Immunosorbent Assay for Detection of Antibodies against African Swine Fever Virus. Pathogens 10.

22. Olival KJ, Cryan PM, Amman BR, Baric RS, Blehert DS, Brook CE, Calisher CH, Castle KT, Coleman JTH, Daszak P, Epstein JH, Field H, Frick WF, Gilbert AT, Hayman DTS, Ip HS, Karesh WB, Johnson CK, Kading RC, Kingston T, Lorch JM, Mendenhall IH, Peel AJ, Phelps KL, Plowright RK, Reeder DM, Reichard JD, Sleeman JM, Streicker DG, Towner JS, Wang L-F. 2020. Possibility for reverse zoonotic transmission of SARS-CoV-2 to free-ranging wildlife: A case study of bats. PLOS Pathogens 16:e1008758.

23. Prince T, Smith SL, Radford AD, Solomon T, Hughes GL, Patterson EI. 2021. SARS-CoV-2 Infections in Animals: Reservoirs for Reverse Zoonosis and Models for Study. Viruses 13:494.

24. Goraichuk IV, Arefiev V, Stegniy BT, Gerilovych AP. 2021. Zoonotic and Reverse Zoonotic Transmissibility of SARS-CoV-2. Virus research 302:198473–198473.

25. Ainsworth M, Andersson M, Auckland K, Baillie JK, Barnes E, Beer S, Beveridge A, Bibi S, Blackwell L, Borak M, Bown A, Brooks T, Burgess-Brown NA, Camara S, Catton M, Chau KK, Christott T, Clutterbuck E, Coker J, Cornall RJ, Cox S, Crawford-Jones D, Crook DW, D’Arcangelo S, Dejnirattsai W, Dequaire JMM, Dimitriadis S, Dingle KE, Doherty G, Dold C, Dong T, Dunachie SJ, Ebner D, Emmenegger M, Espinosa A, Eyre DW, Fairhead R, Fassih S, Feehily C, Felle S, Fernandez-Cid A, Fernandez Mendoza M, Foord TH, Fordwoh T, Fox McKee D, Frater J, Gallardo Sanchez V, Gent N, Georgiou D, Groves CJ, et al. 2020. Performance characteristics of five immunoassays for SARS-CoV-2: a head-to-head benchmark comparison. The Lancet Infectious Diseases 20:1390–1400.

26. Tan CW, Chia WN, Qin X, Liu P, Chen MIC, Tiu C, Hu Z, Chen VC-W, Young BE, Sia WR, Tan Y-J, Foo R, Yi Y, Lye DC, Anderson DE, Wang L-F. 2020. A SARS-CoV-2 surrogate virus neutralization test based on antibody-mediated blockage of ACE2-spike protein–protein interaction. Nature Biotechnology 38:1073–1078.

27. Mehdi F, Chattopadhyay S, Thiruvengadam R, Yadav S, Kumar M, Sinha SK, Goswami S, Kshetrapal P, Wadhwa N, Chandramouli Natchu U, Sopory S, Koundinya Desiraju B, Pandey AK, Das A, Verma N, Sharma N, Sharma P, Bhartia V, Gosain M, Lodha R, Lamminmäki U, Shrivastava T, Bhatnagar S, Batra G. 2021. Development of a Fast SARS-CoV-2 IgG ELISA, Based on Receptor-Binding Domain, and Its Comparative Evaluation Using Temporally Segregated Samples From RT-PCR Positive Individuals. Frontiers in Microbiology 11.

28. Li T, Wang L, Wang H, Li X, Zhang S, Xu Y, Wei W. 2020. Serum SARS-COV-2 Nucleocapsid Protein: A Sensitivity and Specificity Early Diagnostic Marker for SARS-COV-2 Infection. Frontiers in Cellular and Infection Microbiology 10:470.

29. Fenwick C, Croxatto A, Coste Alix T, Pojer F, André C, Pellaton C, Farina A, Campos J, Hacker D, Lau K, Bosch B-J, Gonseth Nussle S, Bochud M, D’Acremont V, Trono D, Greub G, Pantaleo G, Subbarao K. Changes in SARS-CoV-2 Spike versus Nucleoprotein Antibody Responses Impact the Estimates of Infections in Population-Based Seroprevalence Studies. Journal of Virology 95:e01828–20.

30. Mariën J, Ceulemans A, Michiels J, Heyndrickx L, Kerkhof K, Foque N, Widdowson M-A, Mortgat L, Duysburgh E, Desombere I, Jansens H, Van Esbroeck M, Ariën KK. 2021. Evaluating SARS-CoV-2 spike and nucleocapsid proteins as targets for antibody detection in severe and mild COVID-19 cases using a Luminex bead-based assay. Journal of Virological Methods 288:114025.

31. Burbelo PD, Riedo FX, Morishima C, Rawlings S, Smith D, Das S, Strich JR, Chertow DS, Davey RT, Jr., Cohen JI. 2020. Detection of Nucleocapsid Antibody to SARS-CoV-2 is More Sensitive than Antibody to Spike Protein in COVID-19 Patients. medRxiv: the preprint server for health sciences doi:10.1101/2020.04.20.20071423:2020.04.20.20071423.

32. Thudium Rebekka F, Stoico Malene P, Høgdall E, Høgh J, Krarup Henrik B, Larsen Margit AH, Madsen Poul H, Nielsen Susanne D, Ostrowski Sisse R, Palombini A, Rasmussen Daniel B, Foged Niels T, Caliendo Angela M. Early Laboratory Diagnosis of COVID-19 by Antigen Detection in Blood Samples of the SARS-CoV-2 Nucleocapsid Protein. Journal of Clinical Microbiology 59:e01001–21.

33. Le Hingrat Q, Visseaux B, Laouenan C, Tubiana S, Bouadma L, Yazdanpanah Y, Duval X, Burdet C, Ichou H, Damond F, Bertine M, Benmalek N, Choquet C, Timsit J-F, Ghosn J, Charpentier C, Descamps D, Houhou-Fidouh N, Diallo A, Le Mestre S, Mercier N, Paul C, Petrov-Sanchez V, Malvy D, Chirouze C, Andrejak C, Dubos F, Rossignol P, Picone O, Bompart F, Gigante T, Gilg M, Rossignol B, Levy-Marchal C, Beluze M, Hulot JS, Bachelet D, Bhavsar K, Bouadma L, Chair A, Couffignal C, Da Silveira C, Debray MP, Descamps D, Duval X, Eloy P, Esposito-Farese M, Ettalhaoui N, Gault N, Ghosn J, et al. 2021. Detection of SARS-CoV-2 N-antigen in blood during acute COVID-19 provides a sensitive new marker and new testing alternatives. Clinical Microbiology and Infection 27:789.e1–789.e5.

34. Li T, Wang L, Wang H, Li X, Zhang S, Xu Y, Wei W. 2020. Serum SARS-COV-2 Nucleocapsid Protein: A Sensitivity and Specificity Early Diagnostic Marker for SARS-COV-2 Infection. Frontiers in cellular and infection microbiology 10:470–470.

35. Di B, Hao W, Gao Y, Wang M, Wang Y-d, Qiu L-w, Wen K, Zhou D-h, Wu X-w, Lu E-j. 2005. Monoclonal antibody-based antigen capture enzyme-linked immunosorbent assay reveals high sensitivity of the nucleocapsid protein in acute-phase sera of severe acute respiratory syndrome patients. Clinical and Vaccine Immunology 12:135–140.

36. Lu X, Wang L, Sakthivel SK, Whitaker B, Murray J, Kamili S, Lynch B, Malapati L, Burke SA, Harcourt J, Tamin A, Thornburg NJ, Villanueva JM, Lindstrom S. 2020. US CDC Real-Time Reverse Transcription PCR Panel for Detection of Severe Acute Respiratory Syndrome Coronavirus 2. Emerging infectious diseases 26:1654–1665.

37. Li K, Huang B, Wu M, Zhong A, Li L, Cai Y, Wang Z, Wu L, Zhu M, Li J, Wang Z, Wu W, Li W, Bosco B, Gan Z, Qiao Q, Wu J, Wang Q, Wang S, Xia X. 2020. Dynamic changes in anti-SARS-CoV-2 antibodies during SARS-CoV-2 infection and recovery from COVID-19. Nature Communications 11:6044.

38. Wang H, Yuan Y, Xiao M, Chen L, Zhao Y, Haiwei Z, Long P, Zhou Y, Xu X, Lei Y, Bihao W, Diao T, Cai H, Liu L, Shao Z, Wang J, Bai Y, Wang K, Peng M, Liu L, Han S, Mei F, Cai K, Lei Y, Pan A, Wang C, Gong R, Li X, Wu T. 2021. Dynamics of the SARS-CoV-2 antibody response up to 10 months after infection. Cellular & Molecular Immunology 18:1832–1834.

39. Gaebler C, Wang Z, Lorenzi JCC, Muecksch F, Finkin S, Tokuyama M, Cho A, Jankovic M, Schaefer-Babajew D, Oliveira TY, Cipolla M, Viant C, Barnes CO, Bram Y, Breton G, Hägglöf T, Mendoza P, Hurley A, Turroja M, Gordon K, Millard KG, Ramos V, Schmidt F, Weisblum Y, Jha D, Tankelevich M, Martinez-Delgado G, Yee J, Patel R, Dizon J, Unson-O’Brien C, Shimeliovich I, Robbiani DF, Zhao Z, Gazumyan A, Schwartz RE, Hatziioannou T, Bjorkman PJ, Mehandru S, Bieniasz PD, Caskey M, Nussenzweig MC. 2021. Evolution of antibody immunity to SARS-CoV-2. Nature 591:639–644.

40. Peng P, Hu J, Deng H-j, Liu B-z, Fang L, Wang K, Tang N, Huang A-l. 2021. Changes in the humoral immunity response in SARS-CoV-2 convalescent patients over 8 months. Cellular & Molecular Immunology 18:490–491.

41. Mibayashi M, Martínez-Sobrido L, Loo Y-M, Cárdenas WB, Gale M, Jr., García-Sastre A. 2007. Inhibition of retinoic acid-inducible gene I-mediated induction of beta interferon by the NS1 protein of influenza A virus. Journal of virology 81:514–524.

42. Petrovan V, Yuan F, Li Y, Shang P, Murgia MV, Misra S, Rowland RRR, Fang Y. 2019. Development and characterization of monoclonal antibodies against p30 protein of African swine fever virus. Virus Res 269:197632.

43. Martins M, Fernandes Maureen HV, Joshi Lok R, Diel Diego G, Gallagher T. 2022. Age-Related Susceptibility of Ferrets to SARS-CoV-2 Infection. Journal of Virology 96:e01455–21.

44. Palmer Mitchell V, Martins M, Falkenberg S, Buckley A, Caserta Leonardo C, Mitchell Patrick K, Cassmann Eric D, Rollins A, Zylich Nancy C, Renshaw Randall W, Guarino C, Wagner B, Lager K, Diel Diego G, Gallagher T. 2021. Susceptibility of White-Tailed Deer (Odocoileus virginianus) to SARS-CoV-2. Journal of Virology 95:e00083–21.

45. Brown E, Lawson S, Welbon C, Gnanandarajah J, Li J, Murtaugh MP, Nelson EA, Molina RM, Zimmerman JJ, Rowland RR, Fang Y. 2009. Antibody response to porcine reproductive and respiratory syndrome virus (PRRSV) nonstructural proteins and implications for diagnostic detection and differentiation of PRRSV types I and II. Clin Vaccine Immunol 16:628–35.

